# The HIV latency reversal agent HODHBt enhances NK Cell effector and memory-like functions by increasing IL-15 mediated-STAT activation

**DOI:** 10.1101/2021.10.14.464348

**Authors:** Amanda B. Macedo, Callie Levinger, Bryan Nguyen, Mampta Gupta, Conrad Russell Y. Cruz, Katherine B. Chiappinelli, Keith A. Crandall, Alberto Bosque

## Abstract

Elimination of latent HIV reservoirs is a critical endpoint to eradicate HIV. One therapeutic intervention against latent HIV is ‘shock and kill’. This strategy is based on the transcriptional activation of latent HIV with a Latency-Reversing Agent (LRA) with the consequent killing of the reactivated cell by either the cytopathic effect of HIV or the immune system. We have previously found that the small molecule 3-Hydroxy-1,2,3-benzotriazin-4(3H)-one (HODHBt) act as an LRA by increasing Signal Transducers and Activators of Transcription (STAT) activation mediated by IL-15 in cells isolated from aviremic participants. The IL-15 superagonist (N-803) is currently under clinical investigation to eliminate latent reservoirs. IL-15 and N-803 share similar mechanism of action by promoting the activation STATs and have shown some promise in pre-clinical models directed towards HIV eradication. In this work, we evaluated the ability of HODHBt to enhance IL-15 signaling in NK cells and the biological consequences associated with increased STAT activation in NK effector and memory-like functions. We showed that HODHBt increased IL-15-mediated STAT phosphorylation in NK cells, resulting in increased secretion of CXCL10 and IFN-γ and expression of cytotoxic proteins including Granzyme B, Granzyme A, Perforin, Granulysin, FASL and TRAIL. This increased cytotoxic profile results in an increase cytotoxicity against different tumor cell lines and HIV-infected cells. HODHBt also improved the generation of cytokine-induced memory-like NK cells. Overall, our data demonstrate that enhancing the magnitude of IL-15 signaling with HODHBt favors NK cell cytotoxicity and memory-like generation, and targeting this pathway could be further explored for HIV cure interventions.

**Author Summary:** Several clinical trials targeting the HIV latent reservoir with LRAs have been completed. In spite of a lack of clinical benefit of these trials, they have been crucial to elucidate hurdles that ‘shock and kill’ strategies have to overcome to promote an effective reduction of the latent reservoir leading. These hurdles include low reactivation potential mediated by LRAs; the negative influence of some LRAs on the activity of Natural Killer and CD8T effector cells; an increased resistance to apoptosis of latently infected cells; and an exhausted immune system due to chronic inflammation. To that end, finding therapeutic strategies that can overcome some of these challenges could improve the outcome of ‘shock and kill’ strategies aimed towards HIV eradication. In here, we showed that the LRA HODHBt also improves IL-15 mediated NK effector and memory-like functions. As such, pharmacological enhancement of IL-15 mediated STAT activation can open new therapeutic venues towards an HIV cure.

## Introduction

Human immunodeficiency virus (HIV) has caused more than 35 million deaths worldwide. Management of the disease requires the daily administration of a combination of antiretroviral (ART) drugs for the life of the infected individual. This is due to the presence of an intact and inducible latent reservoir of HIV that rebounds after discontinuation of ART therapy [1–3]. Elimination of this latent reservoir is a critical endpoint to eradicate HIV. Therapeutic interventions against latent HIV have been mainly focused on ‘shock and kill’ strategies [4–8]. These strategies are based on the transcriptional activation of latent HIV with a Latency-Reversing Agent (LRA) with the consequent killing of the reactivated cell by either the cytopathic effect of HIV or the immune system. Several clinical trials targeting the latent reservoir with LRAs have been completed [9, 10]. In spite of a lack of clinical benefit of these initial trials, they have been crucial to elucidate hurdles that ‘shock and kill’ strategies have to overcome to promote an effective reduction of the latent reservoir leading to a cure. Among others, these hurdles include (1) low reactivation potential mediated by LRAs through promoting only transcription but low translation of HIV proteins [11]; (2) the negative influence of some LRAs on the activity of Natural Killer (NK) and CD8T effector cells [12–14]; (3) an increased resistance to apoptosis of latently infected cells [15–18]; and (4) an exhausted immune system due to chronic inflammation [19, 20].

Currently, there are three clinical trials involving the IL-15 superagonist N-803 in ART-suppressed people living with HIV (PLWH) (NCT04808908, NCT04340596, NCT04505501). IL-15 is a common gamma chain (γc)-cytokine that promotes its biological effects through activation of the transcription factors Signal Transducers and Activators of Transcription (STAT)-1,3 and 5 [21]. N-803 is the clinical candidate because it has enhanced biologic activity *in vivo* due to longer serum half-life than recombinant IL-15 but both share a common mechanism of action [22, 23]. IL-15 and N-803 have been shown to (1) reactivate latent HIV both *ex vivo* and *in vivo* [13, 24, 25]; (2) enhance NK activity against HIV [26, 27]; (3) improve HIV-specific CD8T cell responses [28]; and (4) promote the migration of NK and CD8T cells to B-cell follicles, a major compartment harboring latently infected cells [29, 30]. However, the clinical benefit of IL-15 or N-803 can be hindered by the transient nature of cytokine signaling. Upon IL-15 binding to its receptor, and activation of the Janus kinase (JAK)/STAT pathway, a series of negative feedback loops are initiated that limit their biological activity and makes the pathway unresponsive to further stimulation [31, 32]. We have recently published that 3-hydroxy-1,2,3-benzotriazin-4(3H)-one (HODHBt) enhanced γc-cytokine signaling in CD4T cells by increasing phosphorylation and transcriptional activity of STATs upon cytokine stimulation [33, 34]. HODHBt retained activated STAT5 in the nucleus and prevented its dephosphorylation and cytoplasm recirculation [33]. HODHBt also enhanced the LRA activity of IL-15 in cells isolated from PLWH [34]. As such, targeting this pathway may enhance the efficacy of using IL-15 or N-803 for cure approaches [29].

IL-15 is also critical for NK cell development, maturation, survival, proliferation, and cytotoxic function [35]. In this work, we evaluated whether targeting IL-15 mediated STAT activation could also improve NK effector and memory-like functions. We demonstrated that HODHBt enhanced IL-15 mediated NK cell activation, as demonstrated by increased expression of activation markers CD25 and CD69; as well as components of cytotoxic cell granules such as Granzyme B, Granzyme A, Perforin and Granulysin, death receptor ligand APO2L/TRAIL and CD95L/FASL expression and promoted higher secretion of IFN-γ and CXCL-10. Moreover, HODHBt enhanced IL-15 mediated cytotoxicity against different tumor cell lines and HIV-infected CD4T cells. Lastly, IL-15, in combination with IL-12 and IL-18, has been shown to confer memory-like properties to NK cells. These properties include a quantitatively and qualitatively increased IFN-γ upon restimulation [36]. We found that addition of HODHBt during the generation of memory-like NK cells led to enhanced IFN-g production upon IL-12 and IL-15 recall.

In conclusion, our results indicate that enhancing cytokine-induced STAT activation with HODHBt, or other small molecules targeting this pathway, may be a suitable pharmacological strategy to enhance NK cell effector and memory-like functions and improve HIV cure strategies.

## Results

### HODHBt enhances IL-15 mediated STAT phosphorylation and transcriptional activity in NK cells

First, we confirmed whether HODHBt enhanced STAT phosphorylation upon cytokine stimulation on NK cells as we have shown before for CD4T cells [33, 34]. For that, NK cells were isolated from peripheral blood mononuclear cells (PBMCs) and treated for 48 hours with DMSO, IL-15, HODHBt or a combination of IL-15 plus HODHBt. After incubation, levels of STAT5, STAT1 and STAT3 phosphorylation were analyzed by western blotting. As expected, IL-15 induced phosphorylation of STAT5, STAT1 and STAT3 over DMSO control (**Figure 1A**). HODHBt alone did not substantially increase the levels of phosphorylation of any of the STATs analyzed. This agrees with our previous studies demonstrating that the activity of HODHBt is dependent upon a γc-cytokine [33]. Combination of IL-15 plus HODHBt induced higher phosphorylation levels of the three STATs than IL-15 alone **(Figure 1A**). Next, we evaluated the transcriptional changes associated with increasing STATs phosphorylation with HODHBt. NK cells were treated overnight with DMSO, IL-15, HODHBt or a combination of IL15 plus HODHBt. Upon incubation, RNA was isolated and subject to RNASeq (**Supplementary Table 1**). IL-15 and the combination of IL-15 plus HODHBt changed the transcription of 2,775 and 4,212 respectively over DMSO control (**Figure 1B** and **Supplementary Table 2**). Interestingly, HODHBt alone did not significantly change the transcription of any gene, in agreement with our previously published data showing minimal activity in the absence of a γc-cytokine (**Figure 1B**) [33]. When comparing the differentially expressed (DE) genes induced by IL-15 and IL-15 plus HODHBt vs DMSO, 89.1% of the genes DE with IL-15 were also DE with IL-15 plus HODHBt. IL-15 plus HODHBt induced an additional 1,738 DE genes compared to IL-15 alone (**Figure 1C**). We then compared the DE genes between IL-15 and IL-15 plus HODHBt. Only 202 were statistically significant, several of which are involved in the effector function of NK cells, including granzyme B (GZMB), granzyme A (GZMA), IFN-γ and CXCL-10 and are well known downstream targets of cytokine signaling (**Figure 1D and Supplementary Table 4**) [37]. Moreover, reactome pathway analysis indicated that the DE genes were involved in cytokine signaling pathways, confirming the specific role of HODHBt enhancing cytokine signaling (**Figure 1E and Supplementary Table 4**).

**Figure 1.**
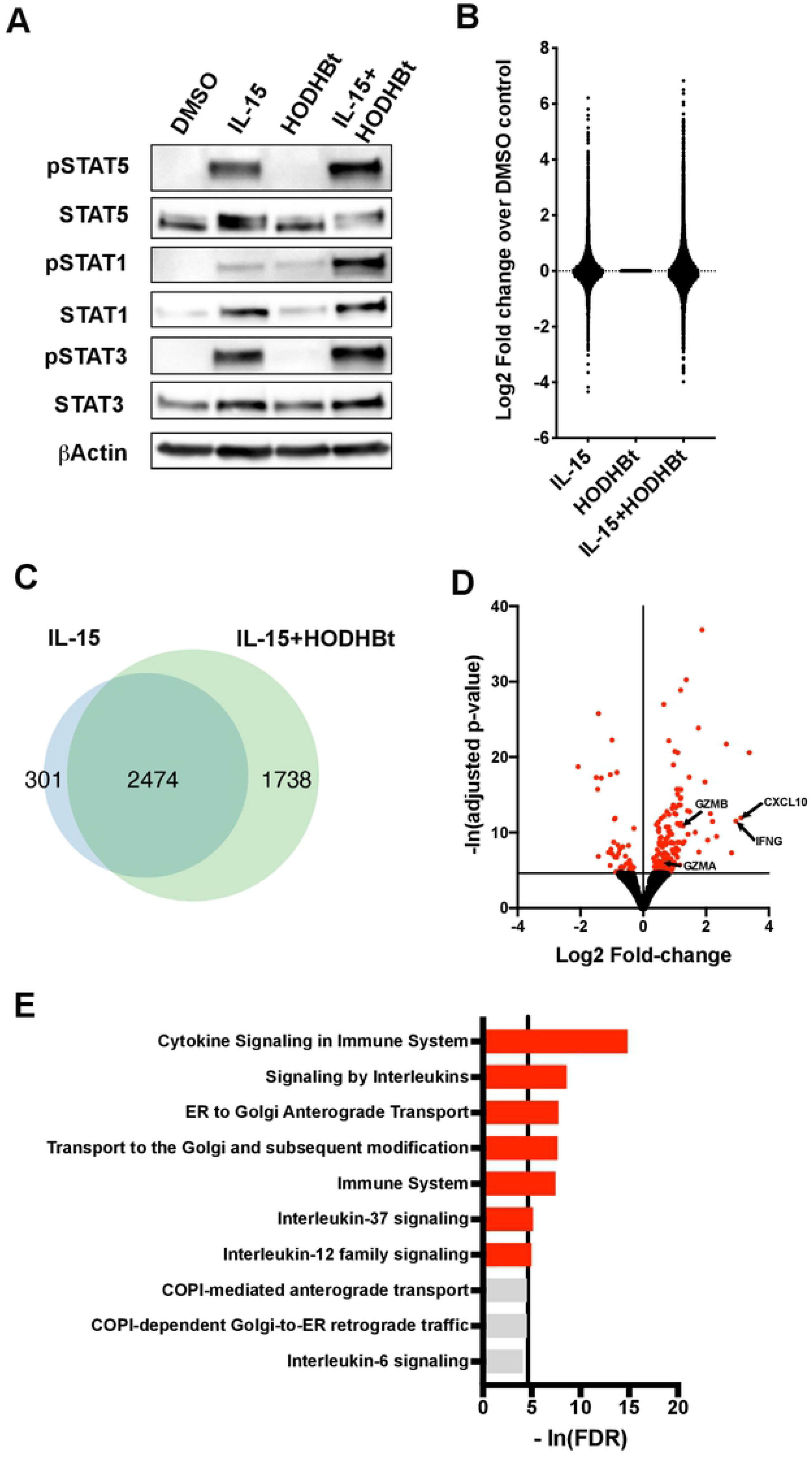
Biological effects of HODHBt in NK cells. **A)** Levels of the phosphorylated and total forms of STAT5, STAT1 and STAT3 in NK cells treated with DMSO, IL-15 (100 ng/ml), HODHBt (100 μM), or a combination of both for 24h. **B)** Log2 fold changes over DMSO control of genes regulated by either IL-15, DMSO or a combination of IL-15 + HODHBt by RNASeq. **C)** Venn-diagram of DE genes between IL-15 vs IL-15/HODHBt over DMSO control. **D)** Volcano plot of DE genes between IL-15 and IL-15 + HODHBt. Genes with adjusted p value <0.01 are indicated in red. **E)** Reactome pathway analysis of DE genes between IL-15 and IL-15 + HODHBt. Pathways with as false discovery rate (FDR) <0.01 are indicated in red.

We then confirmed the induction at the protein level of these and other genes involved in NK effector function in cells isolated from additional 7 to 14 donors (**Figure 2 and Figure 3**). We first evaluated the secretion of the chemokine CXCL-10 and the cytokine IFN-γ by NK cells. The combination of IL-15 plus HODHBt enhanced the secretion of both over IL-15 alone **(Figure 2A-B).** Flow cytometric analysis confirmed enhanced protein expression of Granzyme B and Granzyme A in the IL-15 plus HODHBt condition compared to IL-15 alone (**Figure 2C and D**). Importantly, the presence of HODHBt was not associated with any cell toxicity either alone or in combination with IL-15 (**Figure 3A**). Next, we evaluated the expression of other proteins involved in NK cell cytotoxicity and activation. We observed increased protein expression of Perforin, Granulysin, FASL, TRAIL, as well as the activation markers CD25 and CD69 but not CD16 in IL-15 plus HODHBt compared to IL-15 alone (**Figure 3B-H**). Interestingly, HODHBt alone was sufficient to increase the protein expression of CXCL-10, Granzyme B, Perforin and CD16 over DMSO control (**Figure 2A, 2C, 3B and 3H**). However, this increase was not associated with increased gene transcription based on our RNASeq (**Supplementary Table 1**). Overall, we confirmed that HODHBt enhanced IL-15-induced STAT phosphorylation and transcriptional activity on isolated NK cells leading to increase expression of several cytotoxic molecules important for NK function.

**Figure 2.**
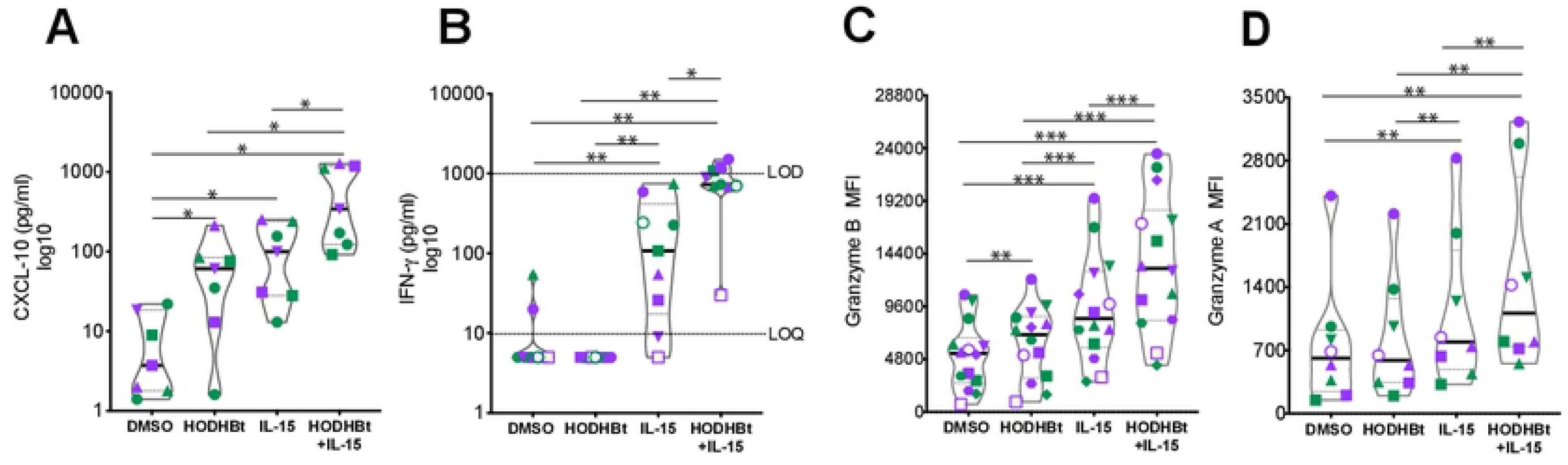
HODHBt enhances secretion of IFN-γ, CXCL-10 in supernatants by NK cells and expression of Granzymes. Concentrations of **(A)** CXCL-10 and **(B)** IFN-γ were quantified in culture supernatants after incubation of NK cells with the indicated treatments for 48h. **(C-D)** Flow cytometric analysis of markers of (**C)** Granzyme B, and Granzyme A after incubation of NK cells with the indicated treatments for 48h. Nonparametric Wilcoxon matched pairs signed rank test was used to calculate p values. Each symbol corresponds to a different donor. Purple symbols are male and green female donors. *P < 0.05, **P < 0.01, and ***P < 0.001, by two-tailed Wilcoxon matched-pairs signed-ranks.

**Figure 3.**
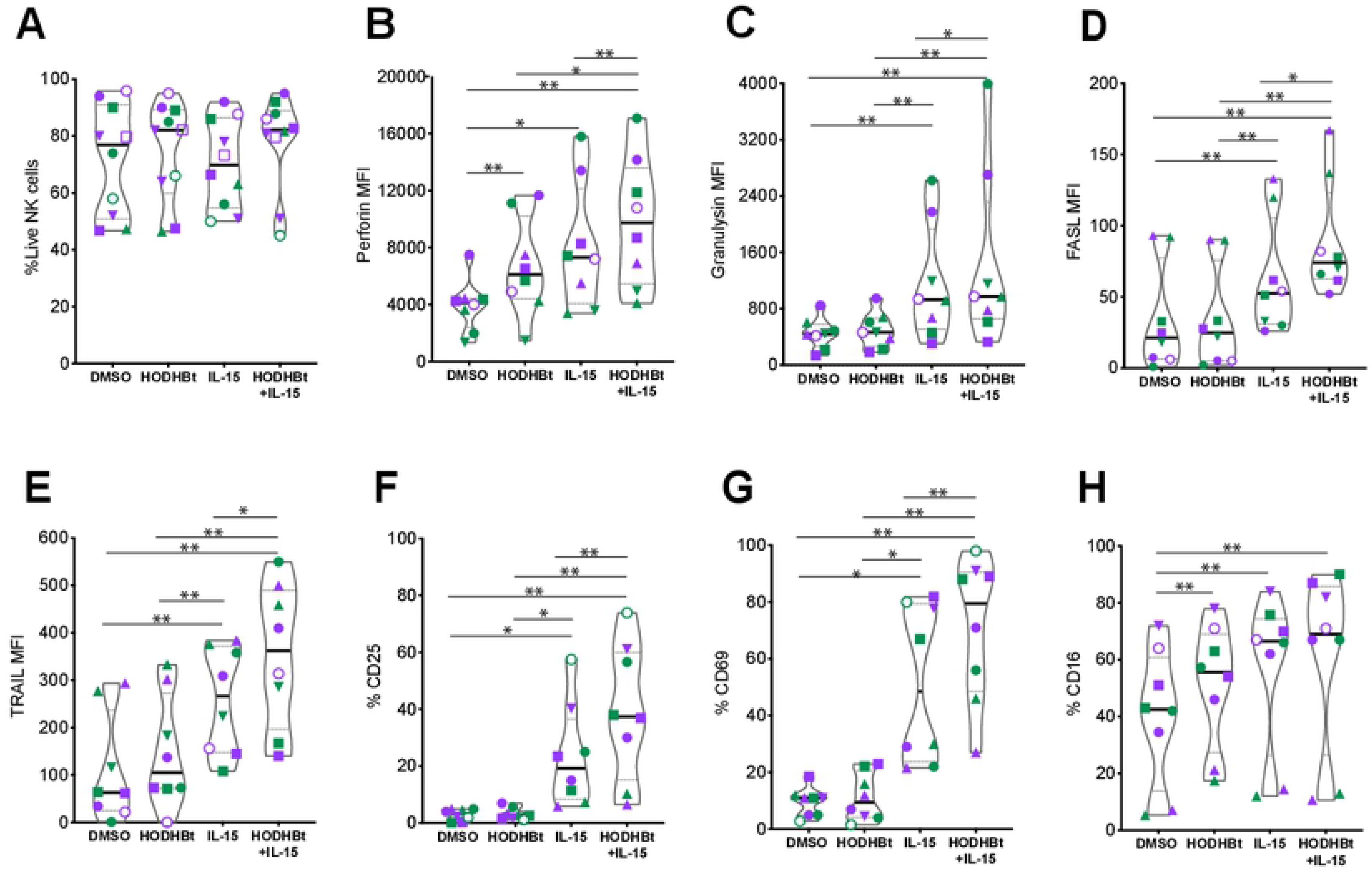
HODHBt enhances the cytotoxic profile of NK cells. Flow cytometric analysis of (**A)** viability; and of markers of cytotoxicity: **(B)** Perforin, **(C)** Granulysin, **(D)** FASL and **(E)** TRAIL; and activation: **(F)** CD25, **(G)** CD69 and **(H)** CD16 after incubation of NK cells with the indicated treatments for 48h. Nonparametric Wilcoxon matched pairs signed rank test was used to calculate p values. Each symbol corresponds to a different donor. Purple symbols are male and green female donors. *P < 0.05, **P < 0.01, and ***P < 0.001, by two-tailed Wilcoxon matched-pairs signed-ranks.

### HODHBt improves the ability of IL-15 activated NK cells to kill tumor cells and HIV-infected cells

Based on the previous results demonstrating that enhancing STAT activation with HODHBt increases the expression of several proteins involved in NK cytotoxicity, we were interested to test whether HODHBt also increases the ability of IL-15-activated NK cells to kill different cancer cell lines, including both hematologic and solid tumors, as well as HIV-infected cells. We first tested whether HODHBt enhanced the ability of IL15-activated NK cells to kill the erythroleukemia K562 cell line, which lacks MHC class I, making these a target for NK cells [38]. NK cells pretreated with either DMSO or HODHBt had low capacity to kill K562 at a 1:1 effector to target (E:T) ratio (**Figure 4A-B**). Pretreatment of NK cells with IL-15 alone induced higher target killing compared to DMSO or HODHBt alone while NK cells treated with the combination of IL-15 plus HODHBt had the highest killing ability of the four groups (**Figure 4A-B**). We then used the bliss independence model to evaluate whether the enhance killing observed with the treatment combination was synergistic [39]. The Bliss model is based on probability theory and assumes that when two drugs act through independent mechanisms, the expected (faxy, e) combinatorial effect should be the sum of the two fractional responses minus their product [(fax+fay)-(fax* fay)]. The interaction of each combination is described by the difference between the observed and the expected response (Δfaxy=faxy, o - faxy,e). Bliss independence analysis yields synergistic (Δfaxy > 0), independent (Δfaxy = 0) or antagonistic (Δfaxy < 0) combinatorial interactions [39]. In this case, the fraction of cell death induced by either HODHBt-treated or IL-15-treated NK cells alone were used to calculate faxy,e and compared with the fraction of cell death induced by NK cells treated with a combination of both (faxy,o). This analysis demonstrates that the combination of IL-15 with HODHBt is synergistic (Δfaxy = 0.05) (**Figure 4B**). The increase NK killing capacity mediated by the combination of IL-15 with HODHBt was also concomitant with an increase in NK cell degranulation and cytokine release, as measured by CD107a expression (**Figure 4C**) and TNF-α production (**Figure 4D**), respectively.

**Figure 4.**
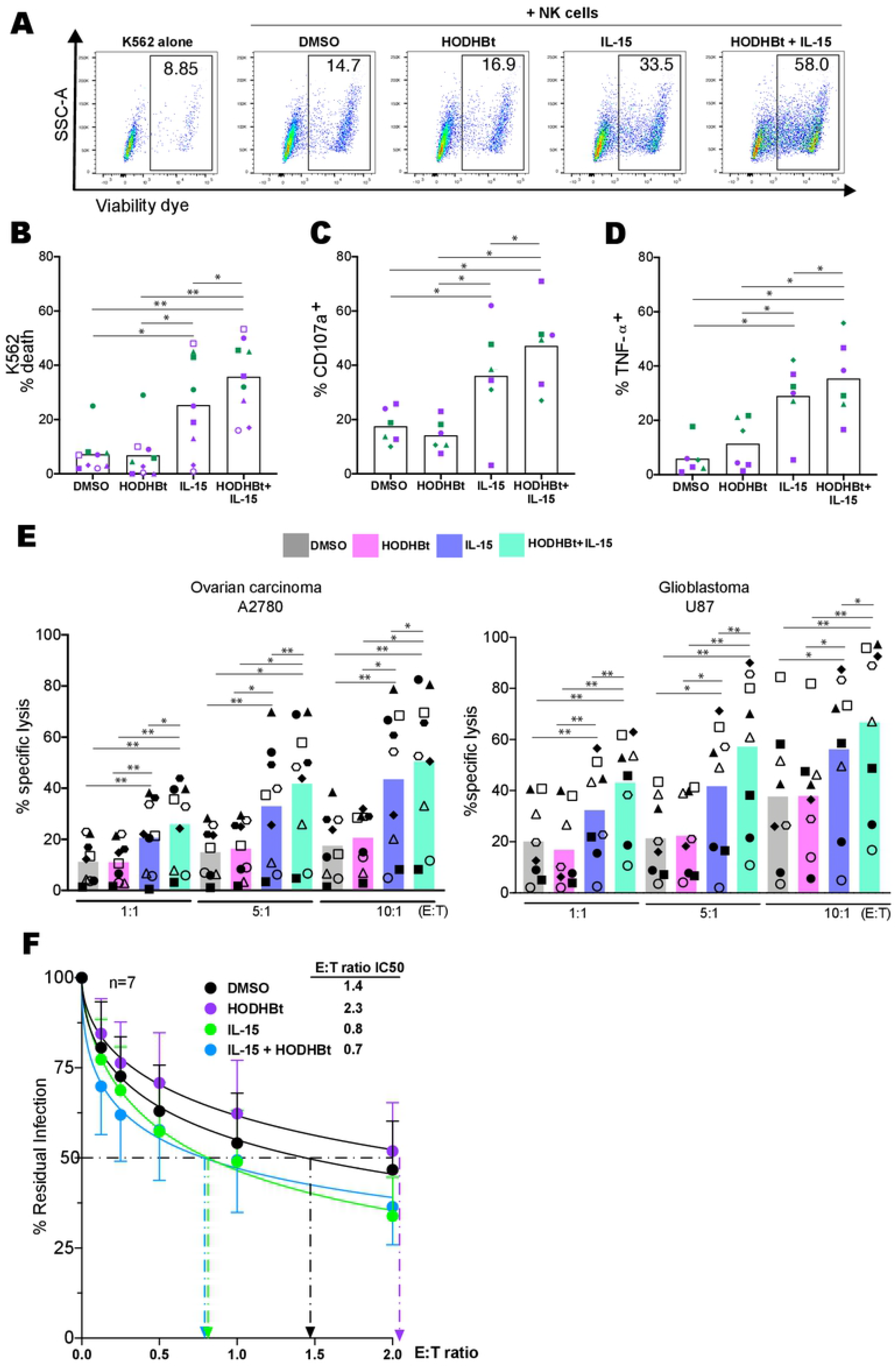
HODHBt enhances IL-15 mediated cytotoxicity of NK cells against cancer cell lines and HIV infected cells. **(A)** Representative flow plots from one donor indicating percentages of cell death of K562 cells cultured alone or in the presence of NK preincubated for 24h at 1:1 Effector to Target (E: T) ratio. **(B)** K562 cell death mediated by pretreated NK cells from 6 male (purple) and 3 female (green) donors. **(C)** Surface expression of CD107a and **(D)** intracellular TNF-α In NK cells co-cultured with K562 cells. **(D)** Percentage of A2780 (left panel) and U87 (right panel) target lysis after incubation with pretreated NK cells at different E:T ratios. **(F)** Percentage of residual HIV-infected CD4T cells in overnight co-cultures of pretreated NK cells and infected CD4T cells at different E:T ratios. IC50 was calculated using nonlinear fit curve. *P < 0.05, **P < 0.01, and ***P < 0.001, by two-tailed Wilcoxon matched-pairs signed-ranks.

We then extended our studies to two models of solid tumors, a human ovarian carcinoma (A2780) and a glioblastoma (U87). Both usually present a bad prognosis and response to current therapies [40]. IL-15 enhanced NK killing of both cell lines at different E:T ratios (**Figure 4E**). As for K562, HODHBt alone did not increase NK cytotoxicity but it did enhance the ability of IL-15-treated NK cells to kill both tumor cell lines (**Figure 4E**). However, the ability of IL-15 or the combination of IL15 plus HODHBt to increase the killing capacity of NK cells is not universal for all cancer types. Neither IL-15 nor the combination increased substantially the ability of NK cells to kill the germinal center OCIly1 or B-cell lymphoma OCILy10 cell lines in comparison to untreated NK cells (**Supplementary Figure 1**).

Finally, we were interested in testing whether HODHBt would increase NK cell killing of HIV-infected CD4 T cells. As HODHBt has been shown to reactivate latent HIV [33, 34], it will be important to address whether targeting this pathway can also enhance the ability of NK cells to kill HIV infected cells. It is known that HIV-infected CD4T cells are naturally resistant to NK cell killing [41–43]. We observed that co-culturing IL-15-activated NK cells with infected autologous CD4T cells induced higher killing of HIV-infected CD4T cells compared to DMSO-treated NK cells, leading to a 42.5% reduction of E:T ratio IC50, similar to previously published by others [26]. The addition of HODBHt to IL-15 modestly enhanced the killing of infected cells over IL-15 alone (12% E:T ratio IC50 reduction over IL-15 alone) (**Figure 4F**). In conclusion, we demonstrate that HODHBt has the ability to enhance the killing capacity of NK cells upon IL-15 stimulation to different cancer cell models and may favor the killing of HIV-infected CD4T cells.

### NK cell phenotype and killing capacity may be affected by age, but not biological sex

We observed a significant amount of variability in the response of NK cells to either IL-15 or the combination of IL-15 plus HODHBt. We decided to evaluate whether biological sex or age of the donors could be a contributor factor to such variability. First, we evaluated granzyme B expression. Although there was no correlation between the baseline expression of granzyme B and age with any of the conditions tested (**Supplementary Figure 2A and Supplementary Table 5**), the induction of granzyme B expression (calculated as fold induction over DMSO control) was significantly associated with age for IL-15 plus HODHBt (p=0.0007, Spearman r=0.81) (**Supplementary Figure 2B**). On the other hand, we did not observe any difference regarding the ability of IL-15 or the combination of IL-15 plus HODHBt to induce expression of granzyme B between female and male donors (**Supplementary Figure 2C**).

Next, we assessed whether age or biological sex influence NK killing capacity. We observed an inverse correlation between the ability of NK cells to kill K562 and the age of the donors in both DMSO or HODHBt treated cells. Both IL-15 and IL15 plus HODHBt can rescue this phenotype (**Supplementary Figure 3A and Supplementary Table 5**). Interestingly, this phenotype seems to be specific for K562 cells, as it was not observed with any of the other cell lines tested (**Supplementary Figure 3B-E**). Finally, there was no difference comparing the killing capacity from NK cells from female and male donors upon any of the stimulations tested (**Supplementary Figure 3F-J**). These data suggest that age, but not biological sex, could be an intrinsic variable that influences the NK cell killing capacity of certain cancer types. IL-15 and the combination of IL-15 plus HODHBt can rescue this phenotype.

### Long-term exposure to HODHBt does not cause exhaustion of NK cells

Higher activation of NK cells with HODHBt could lead to anergy and alteration of NK cell killing ability. To evaluate whether long-term exposure to HODHBt and chronic STAT activation could be detrimental for NK function, we used a commercial medium for NK cell expansion that requires addition of the γc-cytokine IL-2. IL-2, like IL-15, activates STAT5, 3 and 1 and we have shown that HODHBt enhances the activation of STATs mediated by IL-2 [33]. We followed expansion with this protocol for 14 days. The presence of HODHBt through the 14 day-period did not alter the expansion of NK cells or their phenotype based on CD56 and CD16 expression (**Figure 5A** and **5B**). Finally, we evaluated whether long-term exposure of HODHBt alters the killing capacity of NK cells using the erythroblastoma cell line K562 as targets. We did observe that, in general, expansion in the presence of HODHBt increased their killing capacity; albeit it was only statistically significant at the lowest E:T ratio tested (0.0625:1) due to the high variability between donors (**Figure 5C**). These results suggest that long-term exposure of NK cells to HODHBt and continue STAT activation does not induce anergy and may increase their overall killing capacity.

**Figure 5.**
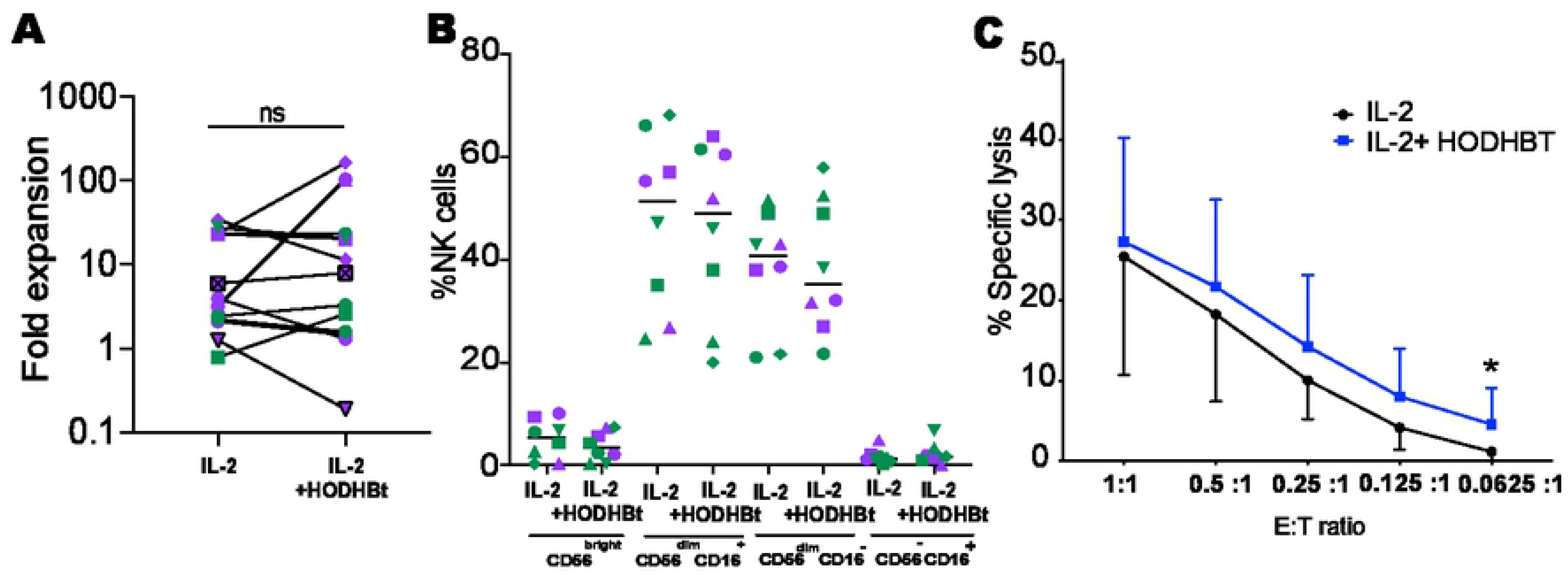
HODHBt does not affect NK cell proliferation rates and phenotype during expansion. **(A)** 2-week fold expansion of NK cells with either IL-2 or IL-2 + HODHBt. **(B)** Percentages of 4 different NK cell subsets based on CD56 and CD16 expression after 14 days of culture. **(C)** Killing of K562 cells at different E:T ratios by expanded NK cells. *P < 0.05 by 2-tailed Wilcoxon matched-pairs signed-ranks was used to compare stimuli.

### HODHBt increases generation of CIML NK cells

NK cells have been shown to have ‘adaptive’ or ‘memory-like’ properties [44–46]. These properties include a quantitatively and qualitatively increase in effector response upon restimulation, characterized by enhanced IFN-γ production [47]. In humans, ‘memory-like’ NK differentiation can be induced by cytokine stimulation with a combination of IL-12, IL-15 and IL-18, and those cells are called cytokine-induced memory-like natural killer cells (CIML) [47, 48]. CIML NK cells are characterized by an epigenetic remodeling of the IFN-γ locus including reduced DNA methylation and enhanced IFN-γ upon restimulation in response to low levels of IL-12 and IL-15, and enhanced survival *in vivo* [47, 48]. Since IL-12 mediates STAT4 activation and IL-15 activates STAT5 [44, 49], we decided to investigate whether HODHBt could increase the generation of CIML NK cells by enhancing STAT activation. We preactivated NK cells using IL-12, IL-15 and IL-18 or control condition in the presence or absence of HODHBt. Next day, cells were washed and cultured in low concentrations of IL-15 without additional HODHBt. After 7 days, NK cells were restimulated with IL-12 and IL-15, and intracellular production of IFN-γ was measured by flow cytometry (**Figure 6A**). As expected, activation with IL-12, IL-15 and IL-18 generated higher levels of CIML NK cells, measured as percentage of cells producing IFN-γ upon cytokine recall, compared to the control group. The addition of HODHBt increased the generation of CIML NK cells for both control and activated conditions (**Figure 6B**). The increase in the generation of CIML NK cells by HODHBt was not associated with appreciable changes in NK phenotype based on CD56 and CD16 expression (**Figure 6C-F**). In conclusion, increasing STAT activation with HODHBt improves the generation of CIML NK cells *in vitro* indicating that the magnitude of STAT activation could be a factor contributing to the generation of memory-like responses in NK cells.

**Figure 6.**
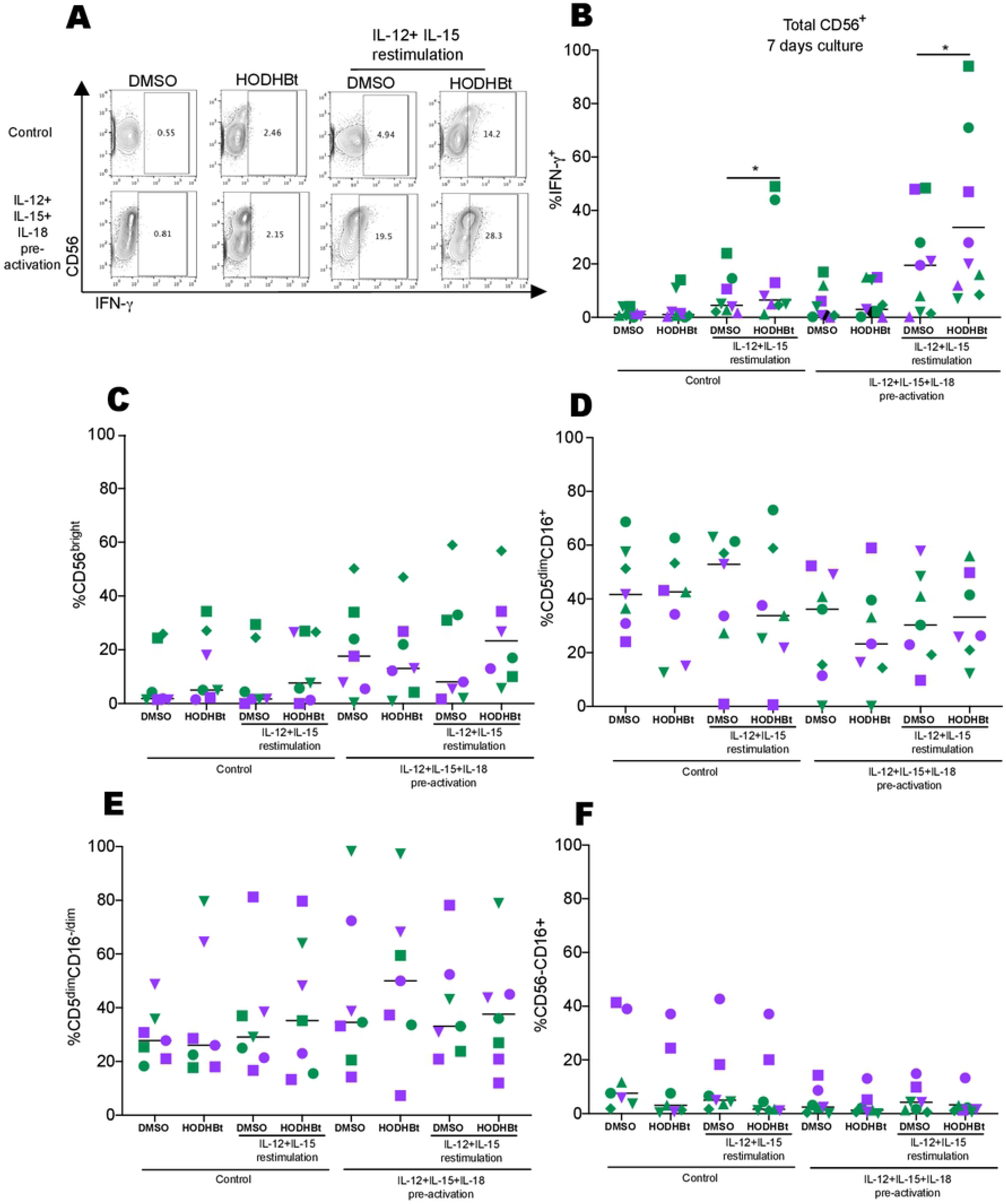
*In vitro* generation of human CIML cells with HODHBt results in enhanced memory response upon recall. **(A)** Representative flow cytometry plots denotating control and pre-activated CIML NK cells expressing IFN-γ with and without recall with IL-12 and IL-15. **(B)** Intracellular IFN-γ after recall in 4 male (purple) and 5 female (green) donors. **(C-F)** Comparisons of NK cell subsets based on CD56 and CD16 expression in 3 male (purple) and 4 female (green) donors. *P < 0.05 by 2-tailed Wilcoxon matched-pairs signed-ranks.

## Discussion

In this study, we tested whether enhancing STAT activation with the LRA HODHBt could enhance cytokine-mediated NK cell cytotoxicity and memory-like generation *in vitro*. We observed that HODHBt increased phosphorylation of STAT5, STAT3 and STAT1 in NK cells treated with IL-15. This increase in phosphorylation by HODHBt was associated with higher STAT transcriptional activation leading to an increased cytotoxic profile phenotype. This was demonstrated by an increased expression of activation markers (CD25 and CD69), cytotoxic proteases (Granzyme A, Granzyme B, perforin, granulysin), death receptor ligands (TRAIL and FASL) and cytokine production (lFN-γ and CXCL-10). Moreover, HODHBt enhanced the ability of IL-15-activated NK cells to kill different tumor cell lines, including chronic myelogenous leukemia, ovarian carcinoma, and glioblastoma, and favored the killing of HIV-infected CD4T cells. Finally, HODHBt also improved the generation of CIML NK cells.

NK cells are part of the innate immune system and play an important role in controlling HIV infection. NK cells are important in the control of SIV replication in the B cell follicles of African Green Monkeys (AGM) [50]. Furthermore, a recent vaccination study using a pentavalent HIV vaccine that builds upon the RV114 study revealed a decrease in acquisition of SHIV due to an increase in the activity of NK cells [51, 52]. Finally, Saez-Cirion et al. showed stronger NK responses in post-treatment controllers from the VISCONTI study than non-controllers [53]. IL-15, through the activation of STATs, is critical for NK cell development, maturation, survival, proliferation, and cytotoxic function [35, 54]. In fact, STAT5 has been proposed as the master transcriptional regulator and plays a role NK cell maturation, survival and cytotoxicity [55–57]. As such, our results suggest that pharmacologic enhancement of IL-15 mediated STAT activation could improve NK activity against HIV. Using an *in vitro* primary model, we observed that IL-15 enhanced the killing of HIV infected cells compared to the controls and its combination with HODHBt slight improved killing capacity. Chronic HIV-1 infection leads to pathologic changes in NK cells, including defective functionality, and control of viremia with ART has been reported to restore some but not all NK cells activity in PLWH [58–61]. Those impairments include lower NK-cell cytotoxicity, IFN-γ production, especially in ART patients with incomplete recovery of CD4 counts [62]. Moreover, HIV-infection even in the context of ART has been associated with higher risk to develop certain types of cancer [63, 64]. As such, identifying pathways that can enhance NK effector function may benefit PLWH not only to control HIV infection but also to reduce the risk for cancer development. For example, robust NK effector functions could lead to better control of HIV replication [51–53]. Furthermore, there are three clinical trials involving the IL-15 superagonist N-803 in ART-suppressed PLWH to promote elimination of latent HIV (NCT04808908, NCT04340596, NCT04505501). Although HODHBt is not currently a clinical candidate, our studies suggest that enhancing IL-15-mediated STAT activation could be a potential mechanism to synergize with N-803 to reactivate latent HIV and rescue NK effector activity. Further efforts will be needed to develop this class of molecules into a clinical candidate or to identify novel compounds that target the same pathway.

Modulation of STAT activation with HODHBt could have an immediate application. The clinical evaluation of NK cells for use in adoptive cell immunotherapies has increased in the last decade [65–70]. NK cells are an alternative to T cell immunotherapies because they preferentially target transformed cells, without the need for prior sensitization or known antigens. Also, NK cells are not thought to elicit graft-versus-host disease and “universal donor” off-the-shelf NK cells are being developed. Despite their promise, chronic exposure to cancer cells leads to impairment of NK cell function. For this reason, multiple strategies have been developed to boost anti-tumor effect of NK cells and abolish tumor resistance. Some examples include adoptive transfer of NK cells after *ex vivo* activation and expansion; restoration of NK cell function using immune checkpoint inhibitors and monoclonal antibody; or cytokine treatment. In this work, we have performed a proof-of-concept study to demonstrate that enhancing IL-15-mediated STAT activation using the small molecule HODHBt can improve NK cell responses. We have observed that HODHBt improved NK mediated killing of several cancer cell lines representative of erythroblastoma, ovarian cancer and glioblastoma compared to treatment with only IL-15 alone. However, we have observed that B cell lines are less sensitive to enhanced NK killing by IL-15 alone or in combination with HODHBt. These data indicate that modulating the STAT pathway may not be sufficient to enhance the killing of some cancer types due to intrinsic cell resistance and additional strategies will be needed. It will be interesting to test whether combination therapy of HODHBt with other current strategies against B-cell lymphomas, such as NK cells expressing chimeric antigen receptors (CAR) against myeloid antigens, could overcome this resistance [71]. Although adoptive NK cell therapy exhibits promise for both cancer and HIV therapy, the development of additional robust methods to expand large numbers of highly effective NK cells is an important area of research. In this study, we tested the effect of HODHBt supplementation in a commercially available NK cell expansion medium and IL-2. The supplementation did not affect the proliferation rates nor the phenotype of the NK cells but increased the ability of NK cells to kill K562. It would be interesting to test whether HODHBt would enhance expansion using different proliferation protocols and other cytokines [72–74].

Although NK cells have traditionally been classified as cells of the innate immune system, they have been demonstrated to have memory features mounting recall responses upon recall [46, 75, 76]. In the context of HIV, pre-existing memory-like NK cells can control viremia in primary infection [77, 78]. Furthermore, SIV-infected macaques but not uninfected macaques have memory-like NK cells able to kill in an antigen-specific manner dendritic cells pulsed with either Gag or Env peptides [79]. A recent study by the Luban group has identified that ongoing viral replication in HIV-infected individuals increases the proportion of memory-like NK cells [80]. In addition, it has been demonstrated that NK cells preactivated *in vitro* with IL-12, IL-15, and IL-18 (so called CIML NK cells) produced higher levels of IFN-γ upon cytokine restimulation and could be detected in higher numbers in comparison to IL-15-preactivated NK [47, 48, 81]. In this work, we were able to demonstrate that HODHBt also enhanced the generation of CIML NK cells based on higher IFN-γ recall response. This suggests that the magnitude of STAT activation could be a contributor factor molding the development of a memory-like phenotype in NK cells. Further studies are warranted to delineate the specific mechanisms involved in this process.

In conclusion, our data show that enhancing cytokine-induced STAT activation with HODHBt significantly increases NK cell cytotoxicity phenotype and function and the generation of CIML NK cells. HODHBt could be further exploited for cell adoptive immunotherapeutic approaches using NK cells against cancer and HIV. Furthermore, the development of clinically relevant compounds targeting this pathway could be an attractive area of research to enhance *in vivo* the effector function of NK cells for HIV cure approaches or against different malignancies and other infectious diseases.

## Materials and Methods

### Cell lines and reagents

Human rIL-2 and IL-15 were provided by the BRB/NCI Preclinical Repository. IL-12 from Peprotech and IL-18 from R&D systems. HODHBt was purchased from AK Scientific. K562-GFP (ATCC CCL-242-GFP) cell culture was maintained in DMEM medium supplemented with 10% fetal bovine serum, glutamine and penicillin-streptomycin. Nelfinavir and Raltegravir were obtained through the AIDS Research and Reference Reagent Program, Division of AIDS, NIAID; and HIV-1NL4-3 from Dr. Malcolm Martin.

### Primary cell culture

Primary NK cells and naïve CD4T cells were isolated from PBMCs by negative selection with cell-type specific EasySep. NK-cell Enrichment Kit (Stem Cell Technologies) according to manufacturer’s protocol. The purity of isolated NK (CD3−CD56+CD16−/+) was verified by flow cytometric analysis. NK cells were stimulated with 100ng/ml IL-15, 100μM HODHBt or IL-15+ HODHBt overnight or at indicated times in the figure legends. For expansion experiments, enriched NK cells were cultured in MACS medium (130-114-429) from Miltenyi Biotech for 14 days (with 500 IU/ML of IL-2 in the absence of presence of 100μM HODHBT) following the manufacture protocol. For the CIML-NK cells experiments, NK cells were plated at 2-5 x10^6^ cells/mL and preactivated for 16 hours using rhIL-12 (10 ng/mL), rhIL-18 (50 ng/mL) and rhIL-15 (1 ng/mL) in the presence of HODHBt or DMSO control and cultured in complete RPMI 1640 medium containing 10% human AB serum (Sigma-Aldrich). As a control, NK cells without any cytokine were plated in medium, in the presence or absence of HODHBt. Next day cells were washed, counted, and replated with rhIL-15 (1 ng/mL) to support survival, with 50% of the medium being replaced every 2-3 days with fresh cytokine. After 7 days, cells were harvested, washed, and restimulated with IL-12 (10 ng/ml) + IL-15 (100 ng/ml) for 6 hours in a 96-well round-bottom plate. Protein transporter inhibitor cocktail (eBioscience) was added after 1 hour and cells were stained for surface NK makers (CD56 and CD16) and intracellular IFN-γ (Cytofix/Cytoperm; BD Biosciences).

### RNAseq analysis

RNA from 5-10 million NK cells treated overnight with indicated conditions was extracted using Rneasy Plus kit (Qiagen). Samples were prepared for Illumina sequencing following the manufacturer’s protocol using the TruSeq® stranded total RNA with rRNA depletion using Ribo Zero TM Human/Mouse/Rat. First-strand synthesis was completed using Superscript TM III (ThermoFisher, cat. 18080044), with an extension temperature of 50°C. Sequencing was performed using a High Output v 2.5 (150 cycles) kit (Illumina, cat. 20024907) on a NextSeq 500, with single indexing. Resulting sequence data were quality controlled using FastQC (<u>bioinformatics.babraham.ac.uk/projects/fastqc/</u>) and MultiQC [82]. Low-quality reads were trimmed using Trimmomatic [83]. Differential gene expression was calculated with DESeq2 [84]. Data has been deposited to NCBISRA: PRJNA753488 (https://dataview.ncbi.nlm.nih.gov/object/PRJNA753488?reviewer=r54l29cmjivfu5tiuk2h7dp6so). Reactome pathway analysis was performed using analysis tools on https://reactome.org [85].

### Western-Blotting

10 μg of protein from each condition was used for Western blotting with a 4-15% Criterion TM TGXTM Precast Midi Protein Gel, 18 well, 30uL (Bio-Rad, Hercules, CA, United States (#5671084). Multiple Western blots were run using the same setup to probe for proteins of similar molecular weight. Proteins were electrophoresed in a Bio-Rad Criterion TM Vertical Electrophoresis Cell at 90-120V for 1-2 hours and then transferred to a PVDF Transfer Membrane (#88518, Bio-Rad, Hercules, CA, United States) using the Bio-Rad Trans-Blot Turbo Transfer System. Membranes were blocked in TBST containing 5% milk or 5% BSA for phosphorylated proteins (blocking buffer) for 1 hour and then incubated in TBST containing 2% milk or 5% BSA (incubation buffer) and primary antibody overnight according to the dilution factor suggested by the manufacturer. The primary antibodies used were pSTAT5 (D47E7, cat 9351S), STAT5 (clone D206Y, cat 94205S), pSTAT3 (clone D3A7, cat 9145P), STAT3 (clone 124H6, cat9139S), pSTAT1 (clone Ser727, cat 9177S), STAT1 (clone 9H2, cat 9176S) (Cell Signaling, Danvers, MA), or the loading control, β-actin (Cat A5441-100UL, Sigma-Aldrich, Saint Louis, MO). Membranes were washed 3 times for 5min each in TBST and incubated in a 1:2000 dilution of HRP-conjugated anti-rabbit or anti-mouse secondary antibody in incubation buffer. Membranes were washed again in TBST and incubated in Immobilon Western Chemiluminescent HRP (Sigma-Aldrich, Burlington, MA, United States (#WBKLS0500) for 5min or 2min for membranes probed with beta actin antibody. Membranes were then imaged and quantified using a GeneGnome imager (Syngene).

### Flow cytometry

To evaluate NK cell activation, cells were washed in PBS then incubated with LIVE/DEAD Fixable Aqua Stain (Invitrogen) for 10 minutes, washed, and the following anti-human antibodies were used for surface staining: CD3-BV786 (clone SP34-2, catalog 563800, BD), CD56-BV605 (clone HCD56, catalog 318334, Biolegend), CD16-FITC (clone 3G8, catalog 555406, BD), CD69-APCCy7 (clone FN50, catalog 310914, Biolegend), CD25-PE (clone BC96, cat 12-0259-42, eBiosciences). To evaluate cytotoxic profile, cells were washed in PBS then incubated with LIVE/DEAD Fixable Aqua Stain (Invitrogen) for 10 minutes, washed, and the following anti-human antibodies were used for surface staining: CD3-BV786 (clone SP34-2, catalog 563800, BD), CD56-BV605 (clone HCD56, catalog 318334, Biolegend). After surface staining, NK cells were fixed and permeabilized with Cytofix/Cytoperm buffer (catalog 554722, BD Biosciences), followed by intracellular staining with Granzyme B-AF700 (clone GB11, catalog 560213, BD Biosciences), Granzyme A-PECy7 (clone CB9, catalog 507221, BD Biosciences), Perforin-FITC (clone dG9, 308104, Biolegend), Granulysin-PE (clone DH2, catalog 308104, Biolegend), TRAIL-APC (clone N2B2, catalog 109310, Biolegend) and FASL-BV421 (clone DX2, catalog 11-0959-42, eBiosciences). For experiments with HIV-infected CD4, cells were stained using the following antibodies: LIVE/DEAD Fixable Aqua Stain (Invitrogen), CD3-BV786 (clone SP34-2, catalog 563800, BD), CD4-APC (S3.5, catalog MHCD0405, BD), CD56-BV605 (clone HCD56, catalog 318334, Biolegend), and intracellular staining with p24/FITC (clone KC57-FITC, catalog 6604665, Beckman Coulter).

Cells were analyzed on a BD LSR Fortessa™ X20 flow cytometer with FACSDIVA™ software (Becton Dickinson, Mountain View, CA) and analyzed using FlowJo (Tree Star Inc., Ashland, OR).

### Cytokine Analysis

Supernatants were collected from each well and stored at −20 °C until ready for analysis with IFN-γ ELISA Kit Invitrogen (cat. # ENEHIFNG2) and CXCL-10 U-PLEX Human IP-10 Assay MSD.

### NK-cell cytotoxicity assays

Flow cytometry-based assays were performed using overnight-activated NK cultures when K562-GFP (ATCC) cells were used as targets. Pretreated NK cells were washed and incubated in duplicates at 1:1 effector-to-target cell (E: T) ratios for 4 h in 96-well U-bottom shaped plates. After incubation, cells were harvested, washed, then stained for flow cytometry. LIVE/DEAD Fixable Aqua Stain (Invitrogen) was used for assessment of dead K562. In parallel cultures, cells were stained to assess NK degranulation and intracellular production of IFN-γ. PE-CD107a (clone H4A) was added to the co-cultures, and after one hour of incubation eBioscience™ Protein Transport Inhibitor Cocktail (eBioscience) was added for an additional 3 hours. After incubation, cells were harvested, washed, stained with LIVE/DEAD Fixable Aqua Stain (Invitrogen), surface-stained with CD56-BV605 (Biolegend), in PBS+3%FBS for 20 min on ice in the dark. Cells were then fixed and permeabilized with Cytofix/Cytoperm buffer (BD Biosciences), washed with Perm/Wash buffer (BD Biosciences), followed by intracellular staining with TNF-α-APC-Cy7 and IFN-γ-PECy7 (Biolegend) for 30 min at 4°C. After washing, cells were resuspended in buffer for cytometric analysis.

DELFIA assay was performed to measure cytotoxicity against cancer cell lines A2780, U87, OCIly1 and OCIl10 according to the manufacturer’s protocol with a few modifications (AD0116, PerkinElmer). Adherent cells lines A2780 and U7 were washed 5 times after labelling with BAPTA. OCIly1 and OCIl10 were washed 3 times after labelling with BAPTA. For experiments using HIV-infected CD4T cells as targets, infected cultured T_CM_ were generated as previously described until day 13, washed and co-cultured with overnight pre-treated NK cells in a 96-well bottom shape plate. Cells were washed and stained for flow cytometry analyses.

### Statistics

Statistical analyses were performed using GraphPad Prism 9.0 software (GraphPad Software). Experiments were analyzed using two-tailed Wilcoxon matched-pairs signed-ranks comparing stimulations and Spearman’s correlation tests. A p value less than 0.05 was considered significant (*P < 0.05, **P < 0.01, ***P < 0.001). All the data with error bars is presented as mean values ± SD.

### Study Approval

Gulf Coast Regional Blood Center - Volunteers 17 years and older served as blood donors. White blood cell concentrates (buffy coat) prepared from a single unit of whole blood by centrifugation were purchased.

## Acknowledgements

Research reported in this publication was supported by the National Institute of Allergy and Infectious Diseases of the National Institutes of Health under Award Number R21/R33 AI116212 and R56 AI145683 to A.B; and an Illumina-GWU Genomics Core Mini-grant. This research has been facilitated by the services and resources provided by District of Columbia, Center for AIDS research, an NIH funded program (AI117970), which is supported by the following NIH co-funding and participating institutes and centers: NIAID, NCI, NICJD, NHLBI, NIDA, NINH, NIA, FIC, NIGGIS and NDDK and OAR. The content is solely the responsibility of the authors and does not necessarily represent the official views of the NIH.

## Authorship contributions

AB and ABM conceived and designed the experiments. ABM and CL performed experiments. MG, CRYC and KBC provided reagents. ABM and AB analyzed the data and wrote the manuscript. BN and KC performed the RNAseq bioinformatic analysis. The authors read and approved the manuscript.

## Disclosure of conflicts of interest

AB and ABM have a patent application on the use of HODHBt to enhance immune responses. The rest of the authors declare no conflict of interest.

